# Mechanical heterogeneity and roles of parallel microtubule arrays in governing meiotic spindle length

**DOI:** 10.1101/385633

**Authors:** Jun Takagi, Yuta Shimamoto

**Affiliations:** Center for Frontier Research, National Institute of Genetics, Yata 1111, Mishima, Shizuoka 411-8540, Japan; Department of Genetics, School of Life Science, SOKENDAI University, Yata 1111, Mishima, Shizuoka 411-8540, Japan

**Author notes:** Correspondence should be addressed to Y.S.

## Abstract

Metaphase spindles are arrays of microtubules whose architecture provides the mechanism for regulated force generation required for proper segregation of chromosomes during cell division. Whereas long-standing models are based on continuous antiparallel microtubule arrays connecting two spindle poles and overlapping at the equator, spindles typically possess a more complex architecture with randomly arranged short filaments. How these heterogeneous multifilament arrays generate and respond to forces has been mysterious, as it has not been possible to directly measure and perturb spindle force while observing relevant filament motility. Here, we combined microneedle-based quantitative micromanipulation with high-resolution microtubule tracking of *Xenopus* egg extract spindles to simultaneously examine the force and individual filament motility *in situ*. We found that the microtubule arrays at the middle of the spindle half are considerably weak and fluid-like, being more adaptable to perturbing forces as compared to those near the pole and the equator. We also found that a force altering spindle length induces filament translocation nearer the spindle pole, where parallel microtubules predominate, while maintaining equatorial antiparallel filaments. Molecular perturbations suggested that the distinct mechanical heterogeneity of the spindle emerges from activities of kinesin-5 and dynein, two key spindle motor proteins. Together, our data establish a link between spindle architecture and mechanics, and highlight the importance of parallel microtubule arrays in maintaining its structural and functional stability.

## Main Text

### Introduction

Spindles are microtubule-based bipolar structures assembled to segregate chromosomes during cell division. Errors in chromosome segregation are linked to aneuploidy, the hallmark of cancer and several developmental disorders in humans (Gordon et al., 2012; Hassold and Hunt, 2001). Forces exerted by the spindle are essential, as they pull chromosomes, monitor erroneous attachment, and control spindle position in a cell (Dumont and Mitchison, 2009b; Inoue and Salmon, 1995). The forces generated in turn act on the spindle and influence its length and bipolarity, which ensure the distance over and axis along which chromosomes are segregated. Therefore, the structure must properly respond to these forces and maintain overall integrity.

Understanding the spindle mechanics requires knowing the internal filament architecture and dynamics, as well as how the arrays of the filaments generate and respond to force. In long-standing models, spindles are described as arrays of long, continuous microtubules radially growing from the opposite spindle poles and forming an antiparallel overlap at the equator (McIntosh et al., 1969; Mitchison and Salmon, 2001; Scholey et al., 2003). Although this relatively simple architecture has been widely observed in small spindles, such as those of yeast (Winey et al., 1995), studies of higher eukaryote spindles, including those of *Caenorhabditis elegans, Xenopus laevis*, and humans, revealed that the filament architecture in these species is much more complex. In particular, the minus-ends of many microtubules are not anchored to spindle poles but instead broadly distributed across the bipolar structure (Burbank et al., 2006; Mastronarde et al., 1993; Redemann et al., 2017). These individual filaments are short and span only part of the spindle, overlapping with each other to form a “tiled-array”-like arrangement (Brugues et al., 2012; Yang et al., 2007). Within this architecture, individual microtubules grow toward either the left or the right spindle pole (Brugues et al., 2012), forming antiparallel as well as parallel filament arrays at varying spindle location. Thus, spindles are architecturally heterogeneous, in terms of filament position and relative filament polarity.

Consistent with this, the poleward flux – the persistent translocation of microtubule lattices characteristic of many metazoans (Ganem and Compton, 2006) – exhibits a non-uniform velocity distribution along the length of the spindle. In particular, the flux speed is substantially faster around the equator (~2-3 μm/min) than nearer the pole (~1 μm/min), which cannot be simply expected from continuous filament lattices spanning from the pole toward past the equator (Burbank et al., 2007; Yang et al., 2008). The poleward movement occurs with filament minus-ends leading (Mitchison, 2005). The dynamics is linked to activities of kinesin-5 and dynein, two key microtubule motors of opposite directionality. *In vitro*, kinesin-5 forms cross-bridges along overlapping microtubules and pushes apart antiparallel filaments (Kapitein et al., 2005). On the other hand, dynein is located at the filament minus-end and may counteract this motion (Tan et al., 2018).

Despite the wealth of information on filament architecture and dynamics, the forces in the spindle are poorly understood. This is because earlier studies have analyzed filament features in a static sample setting, such as one required in electron microscopy, or examined the dynamic samples using live cell methods, but without measuring force. Since Bruce Nicklas’ seminal work (Nicklas, 1997), physical manipulation studies have directly examined spindle forces in cells (Garzon-Coral et al., 2016; Hiramoto and Nakano, 1988; Skibbens and Salmon, 1997); however, the assays used in these studies employed a cell membrane that is robust against physical perturbation and are thus incompatible with molecular tools, or did not accommodate for single filament visualization. Hence, there is little quantitative information on the relationship between spindle force and its architectural dynamics, and thus, the micromechanics of this heterogeneously arranged filament assembly remain mysterious.

Here, to fill this knowledge gap, we combined microneedle-based quantitative micromanipulation with fluorescence speckle microscopy to measure and perturb local spindle force and to simultaneously observe individual microtubule motility *in situ*. The use of *Xenopus* egg extract, which is widely used for studying many cell-cycle events including spindle assembly (Hannak and Heald, 2006), allowed us to perform controlled force perturbation as well as titration over labeled tubulins to achieve conditions necessary for single-filament tracking. We show how the heterogeneously arranged microtubule arrays respond to force and slide apart while maintaining overall stability, and its dependency on motor protein activities.

## Results

### Establishing a method for probing local mechanical responses of microtubules in the metaphase spindle

We used force-calibrated microneedles (stiffness: 0.3–0.5 nN/μm) to probe mechanical responses of microtubules that assemble the *Xenopus* metaphase spindle (Fig. 1A). The microneedles had sufficient bending flexibility and axial rigidity, enabling us to insert the probe tip into the dense microtubule arrays while applying calibrated local force along the pole-to-pole axis of the spindle, the direction along which microtubules roughly align and slide apart. The magnitude of the force applied ranged from 0.4 nN to 1.1 nN, the amount typical for chromosome pulling and spindle positioning in a cell (Garzon-Coral et al., 2016; Zhang and Nicklas, 1999). Microtubule motion response was simultaneously observed using spinning-disk confocal microscope, by tracking the fluorescent ‘speckles’ from X-rhodamine-labeled tubulins (20 nM) incorporated into the filament lattices (probability: <0.4 per filament) (Yang et al., 2007) (Fig. 1B). Images were acquired at ~5 μm above the coverslip surface, a distance that allowed minimal surface friction but sufficient imaging sensitivity.

**Figure 1.**
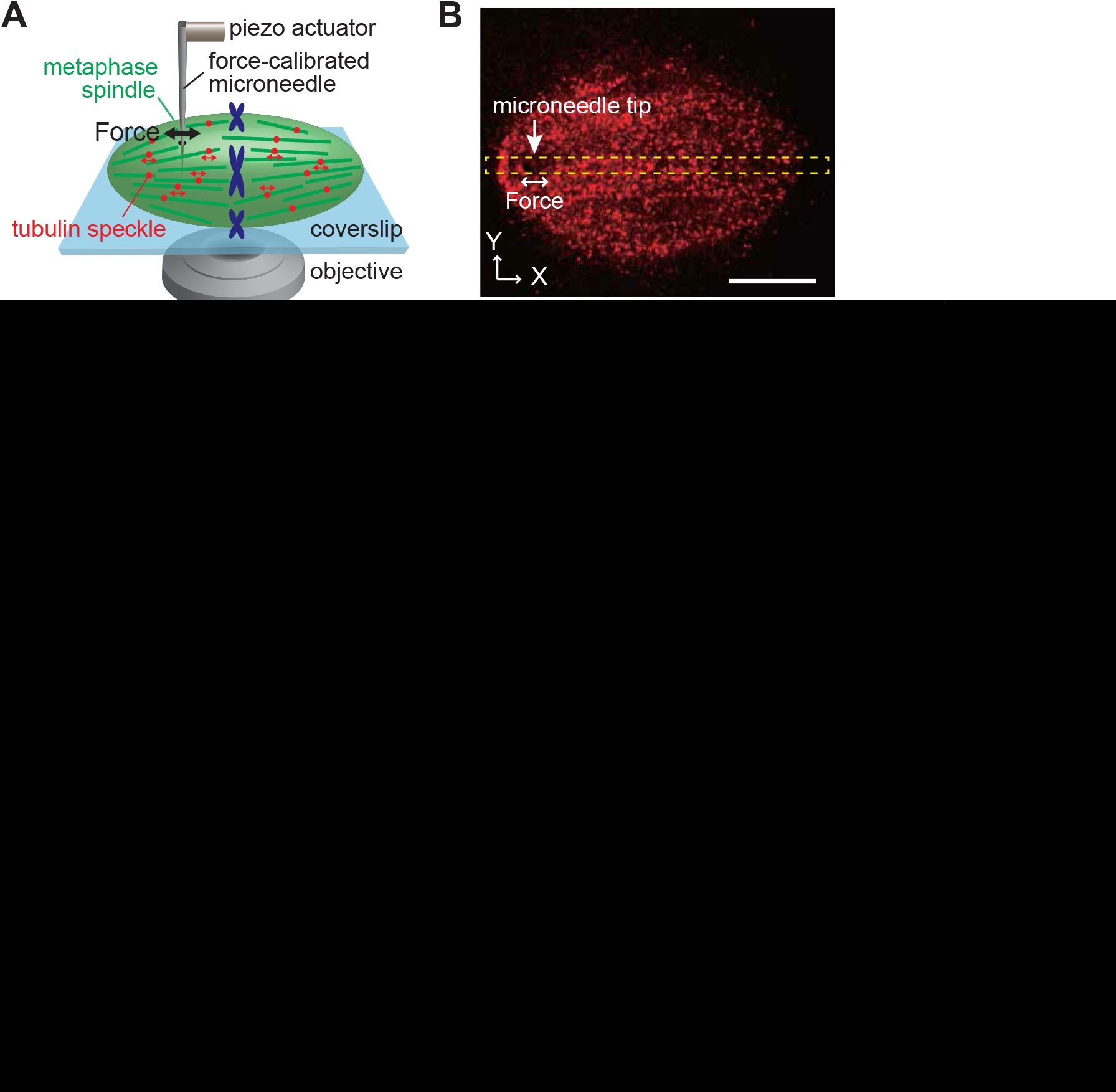
Microneedle-based setup for analyzing *in-situ* spindle microtubule mechanics. (**A**) Schematic of the setup. Single metaphase spindles, assembled in *Xenopus* egg extracts and supplemented with X-rhodamine tubulin (20 nM) for microtubule motion tracking, can be subjected to a calibrated force (0.4–1.1 nN) via the microneedle tip (black double arrow). Microtubule motion responses can be analyzed by tracking fluorescent tubulin “speckles” incorporated into the filament lattices (red dots). (**B**) Representative confocal fluorescence image of a spindle, to which a microneedle tip (arrowhead) was inserted and an external force (double arrow) was applied. Yellow rectangle indicates the region for kymograph analysis. (**C, D**) Kymograph generated along the spindle pole-to-pole axis (**C**) and representative speckle trajectories (**D**), showing the motion of tubulin speckles in the absence of an external force. (**E, F**) An externally applied oscillatory force (frequency: 0.1 Hz; amplitude: 0.2 nN) could entrain this motion while maintaining the overall filament dynamics. Roman numbers above each trace correspond to those in the kymographs. Scale bars, 10 μm (horizontal) and 10 s (vertical). (**G**) Amplitude of speckle motion response upon oscillatory force application was determined by sine-wave fitting of each trajectory and mapped onto a single imaging plane. Warmer color indicates larger response amplitude. Black circle with arrowheads, force application position and direction. (**H, I**) Cross-section of the motion amplitude profile along the long (**H**) and short (**I**) spindle axes. Speckle data within dotted rectangles in (**G**) (width: 5 μm) were projected onto each axis. Blue vertical line, spindle equator position.

Under these conditions and in the absence of an external force, tubulin speckles exhibited persistent poleward translocation at ~2–3 μm/min, while exhibiting stochastic motion fluctuation along the filament’s long axis, in agreeing with previous reports (Yang et al., 2007) (Fig. 1C, D). To determine the mechanical responses of individual microtubules, we needed to eliminate this intrinsic motion ‘noise’. To this end, we developed a method based on oscillatory force input. As shown in Fig. 1E and F, the application of a sinusoidal force at an optimized frequency (0.1 Hz) resulted in periodic back-and-forth movements in the majority of tubulin speckles (>80% of total) without perturbing the overall flux dynamics. The amplitude of the induced speckle movement, which appeared predominantly along the spindle’s pole-to-pole axis (Fig. S1A), was determined based on a least-square fitting to a sinusoidal function (Fig. S1B) and then mapped onto a two-dimensional heat map (Fig. 1G). Projecting the heat map onto the long (Fig. 1H) and short (Fig. 1I) spindle axes revealed the extent of induced speckle movement and its spatial dependency. The speckles’ oscillatory responses did not decay over time (Fig. S1C), provided that our measurements were made while the steady-state metaphase structure was maintained. Further, as the flux dynamics persisted, the direction of the time-averaged speckle movement could be used to define filament polarity. We thus established a method for analyzing local force responses of spindle microtubules depending on filament position and polarity.

### Microtubule arrays at the middle of the spindle half are more mechanically compliant than those near the pole and the equator

Using this approach, we first asked whether the mechanical responses of microtubules differ depending on their position in the spindle (Fig. 2). The microneedle tip was inserted into either of three spindle regions: near the pole, near the equator, or at the middle of the spindle half. Speckle motion amplitude profiles (as in Fig. 1G) were obtained for multiple spindle samples, pooled, and averaged to generate an average motion amplitude profile along the long (Fig. 2A–C) and short (Fig. 2E–G) spindle axes. We found that for forces applied near the spindle pole (<5 μm from the structure’s edge), the profile showed a plateau phase over ±5 μm from the point of force application (<10 % drop), indicating a coupled lattice movement at the vicinity of the spindle pole (Fig. 2A). This concerted lattice movement was also observed when force was applied near the spindle equator (<5 μm from the structure’s center), albeit that the amplitude was maintained toward both spindle poles (Fig. 2C). Notably, however, when force was applied at the middle of the spindle half (i.e. between the pole and the equatorial regions), we observed much steeper amplitude decay (>50% drop) within the same ±5-μm distance (Fig. 2B). Normalized motion amplitude profiles further revealed that the induced relative lattice movement was >2-fold larger at the middle of the spindle half (slope: 0.13 ± 0.05) than near the pole (0.03 ± 0.02) and the equator (0.06 ± 0.05) (n = 5 each, Fig. 2D). Speckles located along the short spindle axis also exhibited substantial movement parallel to the direction of force application (Fig. 2E–H), indicating lateral mechanical coupling between neighboring filaments.

**Figure 2.**
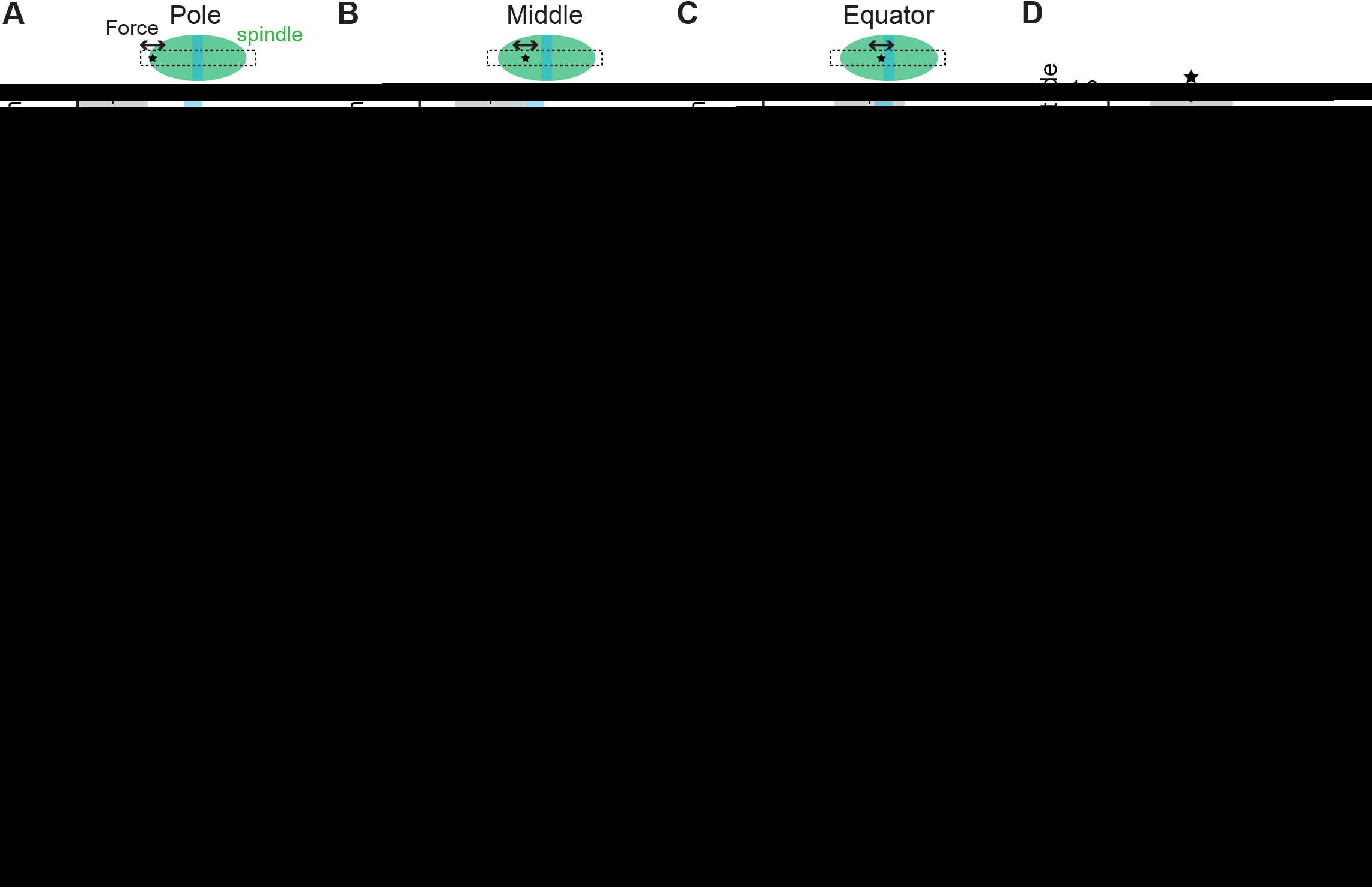
Microtubule arrays at the middle of the spindle half are less mechanically resistant to force than those near the pole and the equator. (**A–C**) Magnitude of speckle motion response depending on position along the long spindle axis, examined using an oscillatory force (0.1 Hz) applied near the spindle pole (**A**), at the middle of the spindle half (**B**), and near the equator (**C**). Data from multiple spindles (grey lines, n = 5 each) were each aligned at baseline, pooled and averaged at 1 μm bin width (navy plots). Bars are S.D. Vertical bars in light blue, approximate equatorial position. (**D**) The averaged profiles in (**A–C**) were each normalized to the peak value and overlaid at the peak position. (**E–G**) The data in (**A–C**) were analyzed for speckles that located along the short spindle axis.(**H**) Normalized amplitude profiles of (**E–G**) generated as in (**D**). (**I**) Local effective stiffness at each spindle region, as estimated on the basis of each motion amplitude profile within ±5 μm from the peak (grey highlighted area in (**A–C**) and (**E–G**)). Data are mean ± SD (n = 5 each). \**p* <0.05 \*\**p*<0.01, two-tailed Student’s *t*-test. N.S., not significant. (**J, K**) Local mechanical properties of the spindle, measured by microrheology (see Fig. S2). Oscillatory forces were applied along the long spindle axis and at varied frequency (0.01–4 Hz) (n = 10 at each spindle location). Dynamic stiffness (**J**) represents total mechanical resistance associated with viscous and elastic deformations. Phase shift (**K**) is a measure of how elastically (*θ* = 0) or viscously (*θ*= π/2 rad) the structure is deformed.

Therefore, the arrays of microtubules are mechanically coupled in both longitudinal and lateral directions, and the coupling strength varies depending on spindle location.

Based on the extent of local lattice movement and the amount of force applied, we estimated the dynamic modulus of the spindle, a measure of the structure’s stiffness (see Materials and Methods). The moduli for the long axis were 5.6 ± 2.0, 1.4 ± 0.2, and 3.2 ± 1.4 kPa (or ×10_3_ pN/μm_2_) at the pole, the middle of the spindle half, and the equator, respectively (magenta, Fig. 2I) (n = 5). On the other hand, the moduli for the short axis were much smaller overall, but also depended on spindle location: 0.7 ± 0.3, 0.3 ± 0.1, and 0.4 ± 0.2 kPa, in the same location order (cyan, Fig. 2I). The values are in an order of magnitude comparable to previously measured macroscopic spindle stiffness (several kPa) (Itabashi et al., 2009; Takagi et al., 2014) and indicate the substantial local mechanical compliance of filament arrays at the middle of the spindle half.

Our previous study showed that the spindle is a viscoelastic material (Shimamoto et al., 2011), and thus, the local mechanical responses may vary depending on the timescale at which forces are applied. As our method allowed for analyzing speckle motion at relatively slow timescales (e.g. 0.1 Hz), we conducted an independent stiffness measurement based on microrheology analysis (Fig. S2A–D). The analysis revealed greater mechanical compliance at the middle of the spindle half than near the pole and the equator, over a range of timescales from minutes to sub-seconds (frequency: 0.02–4 Hz) (Fig. 2J). Moreover, the structure underwent a predominantly viscous, fluid-like deformation at the middle of the spindle half, whereas the structure close to the spindle pole exhibited less fluidity (Fig. 2K). Together, microtubule arrays at the middle of the spindle half engage in a relatively weak, viscous mechanical coupling, whereas those around the spindle pole are more rigid and elastic.

### Spindle microtubule response to force is independent of filament polarity

Microtubules in the spindle orient toward either the left or the right spindle pole, and the proportion varies depending on their location in the spindle. To test whether this filament feature leads to different mechanical outputs, we analyzed the dependency of the speckles’ force responses on filament polarity (Fig. 3). Tubulin speckles were classified into two groups based on the directionality of their persistent poleward movements, and then the force-induced motion amplitude was determined as described above (Fig. 3A). Consistent with previous studies, the two motile fractions appeared nearly equal at the equator (~50:50), while those moving toward the proximal spindle pole became predominant nearer the pole (~80:20), suggesting the predominance of parallel filaments with their minus-ends facing outward (n = 4, Fig. 3B). However, we found no significant differences in speckle motion amplitude depending on the assigned filament polarity, within the accuracy that could resolve its regional variation (Fig. 3C). The analysis was conducted for multiple spindle samples (n = 4), with consistent results (Fig. 3D). These findings suggest that at each spindle location, microtubules engage in approximately equal mechanical coupling regardless of filament orientation. In addition, the compliant filament array at the middle of the spindle half is predominantly parallel.

**Figure 3.**
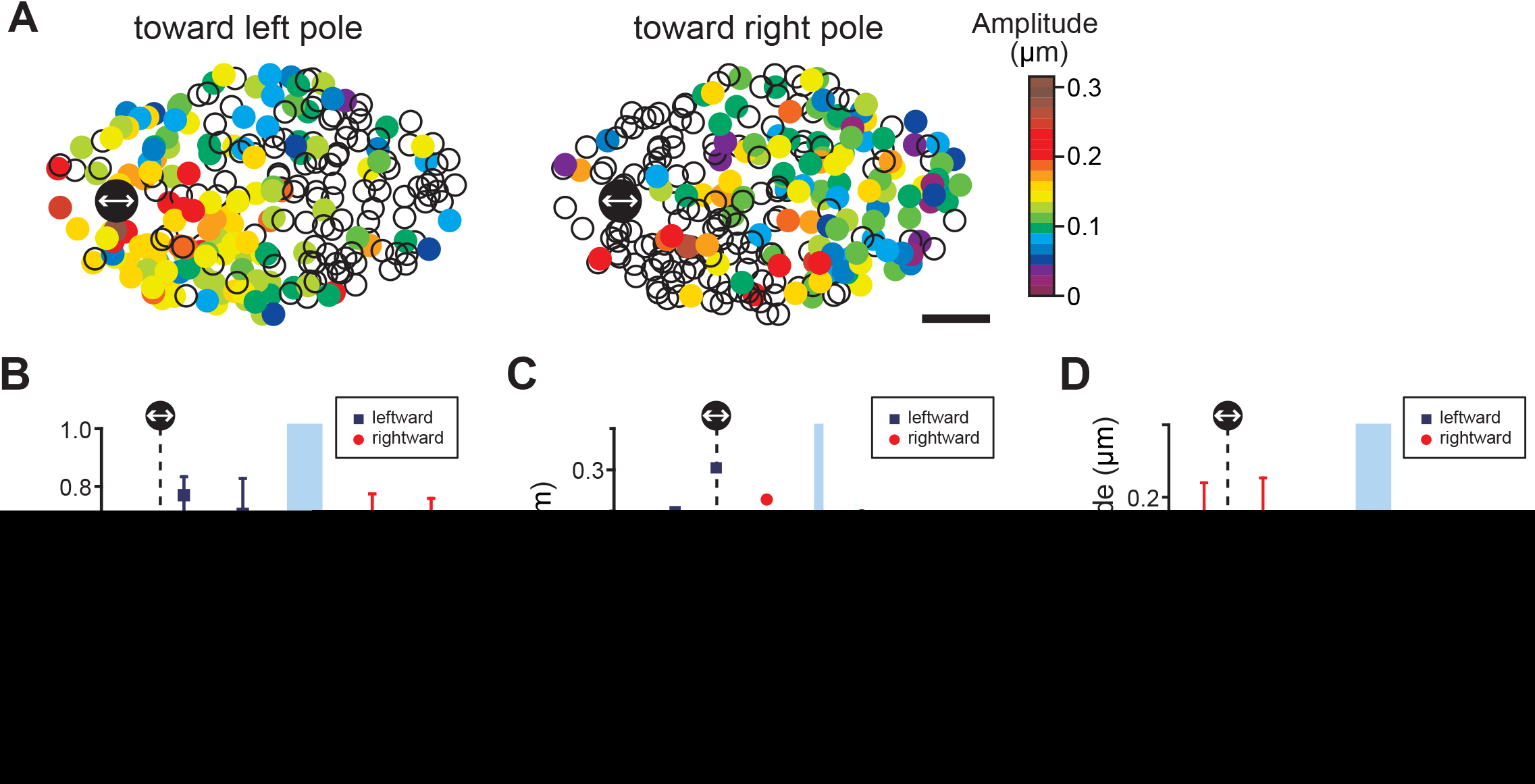
Local mechanical responses of spindle microtubules are independent of filament polarity. (**A**) Two-dimensional heat maps showing the dependency of speckle motion amplitude on microtubule polarity. Speckles that had been moving toward the left and right spindle poles (tracked over >10 s) are analyzed for directionality and mapped in separate panels. Black circle, force application location (frequency: 0.1 Hz; amplitude: 0.25 nN). Warmer color indicates larger amplitude response. Open circles in each map are the fraction of speckles that were assigned the opposite polarity. (**B**) Ratio of leftward-versus rightward-moving speckles (black squares and red circles, respectively) as a function of the position along the long spindle axis. Data from n = 4 spindle samples were pooled and averaged at each 5-μm bin. Bars are SDs. (**C**) Individual speckle motion amplitude in (**A** is projected along the long spindle axis. Speckles moving toward the left and right spindle poles are plotted in different symbols (blue squares and red circles, respectively). Other marks are as in Fig. 1. (**D**) Averaged motion amplitude of leftward- and rightward-moving speckles at different spindle location, obtained from n = 4 spindles. Data as in (**C**) are pooled and averaged at each 5-μm bin. Bars are SDs.

### Microtubule arrays nearer the spindle pole predominantly slide outward against pole-separating force while the dynamics of equatorial filaments are maintained

The local microtubule responses we characterized thus far occurred at sub-micron length scale, while spindles maintained a steady pole-to-pole length. To examine how the filaments respond to force that influences macroscopic spindle-length, we employed a dual-microneedle setup (Takagi et al., 2014) and induced a global length perturbation of the spindle (~20% increase from the steady-state size) (Fig. 4A). Spindles were double-labeled with Alexa 488-tubulin (400 nM) and X-rhodamine-tubulin (20 nM) for imaging their overall morphology and individual microtubule motion dynamics, respectively, and were stretched by moving one microneedle away from the other (Fig. 4B). The stretching speed (100 nm/s) was such that it led to the development of nN-order force across the length of the bipolar structure (Takagi et al., 2014). The analysis was conducted for speckles that could be tracked for >10 s, a period that covers average tubulin turnover in spindles (~30–60 s) (Needleman et al., 2010; Salmon et al., 1984). As shown in the kymograph (Fig. 4C), soon after the microneedle movement was initiated (t = 0), the spindle first underwent a brief period of parallel translocation due to mechanical compliance between the probe tip and the spindle (labeled “Trans” in Fig. 4C), and then continuously elongated until the microneedle motion was stopped (labeled “Stretch” in Fig. 4C). Quantitative analysis revealed that during the course of the stretch, spindle length increased at a nearly constant velocity (96 ± 26 nm/s, n = 4) and reached ~110–130% of the initial pole-to-pole distance (34.2 ± 6.0 μm to 40.7 ± 6.2 μm, n = 4) (orange highlighted area in Fig. 4D). Associated with this change, tubulin speckles moved predominantly parallel to the force application direction and yielded trajectories of various contour lengths (Fig. 4E). Average instantaneous velocities of the speckles were calculated along individual trajectories and then corrected for velocities relative to the spindle center to compensate overall bias toward the moving spindle pole.

**Figure 4.**
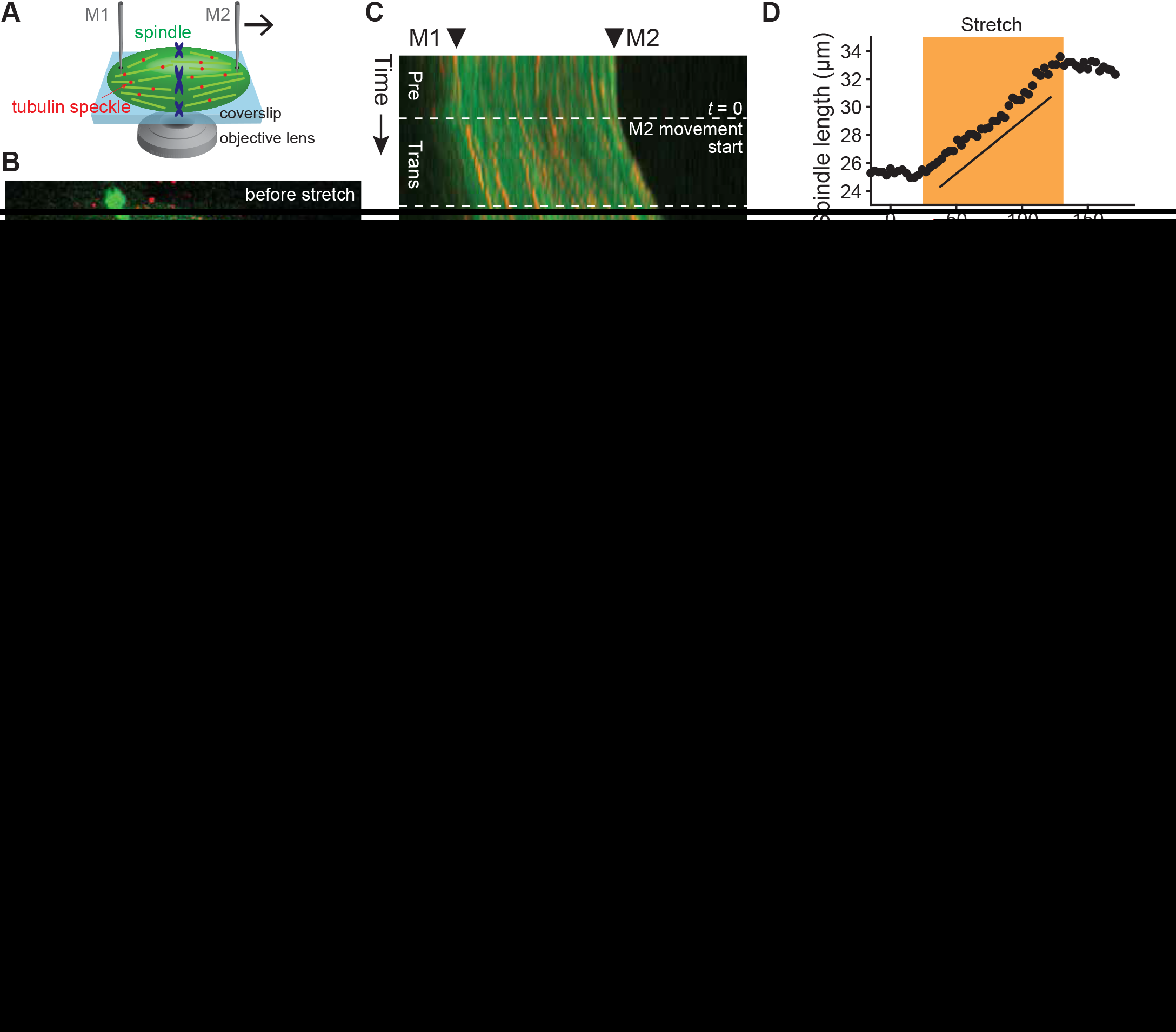
Spindle length change is coupled with sliding of microtubule arrays nearer the pole. (**A**) Dual microneedle-based setup for examining microtubule motion dynamics associated with spindle length change. Microtubules were double-labeled with X-rhodamine-tubulin (red, 20 nM) and Alexa 488-tubulin (green, 400 nM) for speckle imaging and spindle length measurement, respectively. One microneedle (M1) is used to pin down the spindle while the other (M2) can be moved at 100 nm/s to apply outward stretching force. (**B**) Confocal snapshots of a spindle before and during the course of a stretch. Merged images of labeled tubulins (red: X-rhodamine; green: Alexa 488) are shown. Chromosomes were also labeled with SYTOX green dye. Dotted lines indicate changes in microneedle tip positions. Scale bar, 10 μm. (**C**) Kymograph generated along the spindle pole-to-pole axis in (**B**)) (line width: 3 μm). Arrowheads, initial microneedle tip positions. Following the onset of microneedle movement (*t* = 0), the spindle first underwent a brief period of parallel translocation (0–25 s, labeled “Trans”) and was then stretched at a nearly constant velocity (25–135 s, labeled “Stretch”). Horizontal scale bar, 10 μm; vertical scale bar, 20 s. (**D**) Time course of spindle-length change. Orange highlighted area indicates the period over which the spindle was stretched. Slope is 94 nm/s (*R^2^* = 0.98) by linear regression. The following analyses were performed at the highlighted steady stretching phase. (**E**) Speckle motion response. Trajectories of individual speckles that could be tracked for >10 s were projected onto a single imaging plane. Grey ovals with broken and solid lines are approximate spindle positions at the onset and the end of stretch, respectively. (**F–I**) Speckle velocity analyzed before stretch (**F, G**) and during steady stretching phase (**H, I**). Heat maps (**F, H**) were generated on the basis of the average velocity of individual tubulin speckles relative to spindle center and plotted at their initial position along the spindle axes. (**G, I**) Dependence of speckle velocity on the position along the long spindle axis. Histograms indicate overall velocity distribution of all the speckles analyzed. (**J**) Magnitude of the increase in average speckle velocity upon application of a stretching force as a function of the long-axis spindle position. Data obtained from n = 4 spindles were pooled and averaged at every 0.1 relative spindle position bin. Grey lines are trends predicted from a simple multifilament array model (**K**) (see also Fig. S3C), which assumes that the predominant relative filament movement occurred evenly across the spindle (broken line), near the pole (solid line), or near the equator (dotted line).

In the absence of an external force, we observed that the speckles exhibited a non-uniform velocity distribution along the long spindle axis (Fig. 4F, G), consistent with previous reports (Burbank et al., 2007; Yang et al., 2008). The average absolute velocity was 2.4 ± 1.3 μm/min around the equator (n = 135 tracks; blue highlighted area, Fig. 4G) and 1.4 ± 1.0 μm/min nearer the pole (n = 63 tracks; white area, Fig. 4G). When the outward stretching force was applied (Fig. 4H, I), the speed of speckle movement nearer the spindle pole increased to a level that nearly matched the rate of spindle elongation (2.3 ± 1.4 μm/min; n = 33 tracks) (white area, Fig. 4I). Notably, however, this force did not significantly influence the dynamics of speckles located around the equator, where the bidirectional antiparallel movement was maintained at nearly the intrinsic velocity (2.4 ± 1.2 μm/min; n = 93 tracks, blue highlighted area in Fig. 4I). During the course of the stretch, the overall distribution of speckle velocity maintained symmetry, indicating that the manipulation was applied evenly (histograms in Fig. 4G, I). We repeated the analysis for three additional spindles that had been successfully stretched, and consistently observed the preferential acceleration of speckle translocation nearer the pole versus the equator (Fig. S3A). The enhanced speckle motility was most likely caused by induced relative filament sliding, not breakage of the filaments, because a majority of speckles maintained steady translocation speed during the course of stretch (Fig. S3B). Further, the magnitude of the force applied (<1 kPa) was orders of magnitude lower than the tensile strength of microtubules (>1 MPa) (Peter and Mofrad, 2012).

Fig. 4J summarizes the effect of an applied stretching force on the speed of speckle movement at various spindle locations. The profile should yield a straight line if the change in speckle movement is uniform across the spindle length (grey broken line in Fig. 4J; for schematic, see Fig. 4K). On the other hand, the profile deviates from the linear relationship when predominant sliding occurs nearer the pole or the equator (solid and dotted lines, respectively, in Fig. 4J; Fig. S3C). Our data is more consistent with a concave shape, indicating that major filament sliding took place nearer the pole, including the middle of the spindle half (red plots in Fig. 4J). Together, the microtubule arrays nearer the spindle pole adapt to a force that perturbs the spindle’s pole-to-pole distance, while the dynamics of equatorial filament arrays are largely unperturbed.

### Kinesin-5 contributes to the rigidity of microtubule arrays near the pole and the equator

To explore the molecular mechanisms underlying the local microtubule responses, we molecularly perturbed the key spindle motors, kinesin-5 and dynein (Fig. 5). The single microneedle setup was used for these analyses as it enabled us to measure the mechanics of essentially any spindle phenotype, including spindles with fragile poles. Our primary focus was on kinesin-5, which localizes all along the spindle and is enriched near the pole (Sawin et al., 1992). We first used AMPPNP (1.5 mM), a slow-hydrolyzing ATP analogue that immobilizes kinesin-5 onto the microtubule lattice in the “rigor” state (Kapoor and Mitchison, 2001). Dynein is relatively insensitive to AMPPNP (Heald et al., 1996). This treatment did not significantly alter overall spindle length and bipolarity (Fig. 5A); however, we found a global reduction in mechanical responses of the microtubule arrays across the length of the bipolar structure (Fig. 5B–D, Fig. S4A). The local dynamic modulus (stiffness) increased accordingly (11.7 ± 2.7, 3.5 ± 1.6, and 4.0 ± 1.6 kPa at the pole, middle, and equator; n = 3 each), with statistical significance at the pole and the middle of the spindle half (Fig. 5E), suggesting that persistent cross-bridges were made between overlapping microtubules.

**Figure 5.**
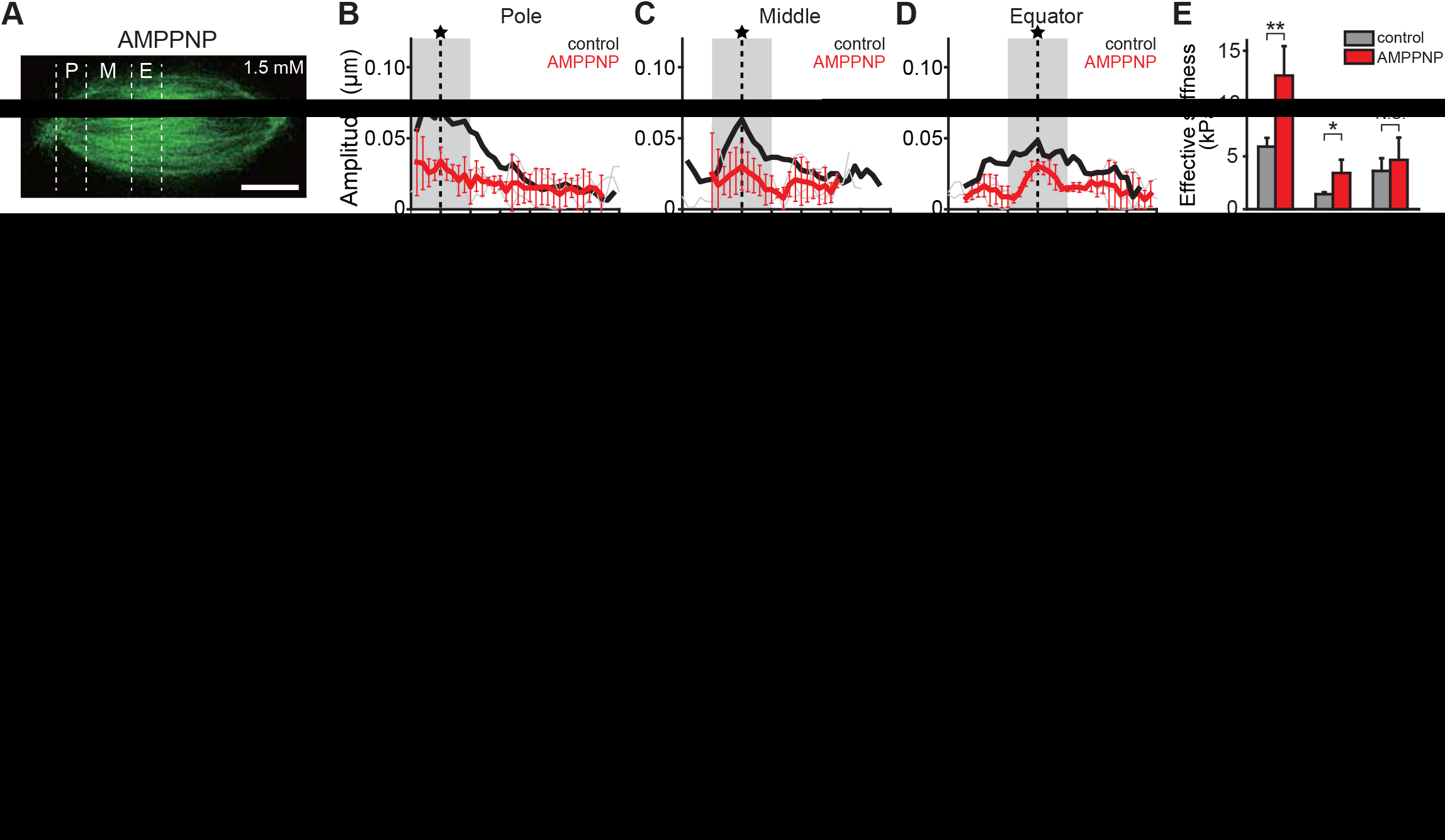
Effect of motor protein inhibition on the local mechanical responses of microtubules. Results of molecular perturbation assays with 1.5 mM AMPPNP (n = 3) (**A–E**), 10 μM or 20 μM monastrol (n = 3 and 5) (**F–J**), and 1 mg/ml anti-dynein 70.1 antibody (n = 4) (**K–O**). (**A, F, K**) Representative confocal snapshots of spindles upon drug treatment. Local microtubule responses were measured using an oscillatory force (0.1 Hz) at various spindle locations (indicated by white dotted lines). Average motion amplitude profiles were then generated along the long spindle axis measured near the spindle pole (**B, G, L**), at the middle of the spindle half (**C, H, M**), and at the equator (**D, I, N**). Averaged profiles from non-treated spindles are shown for comparison (black, corresponding to Fig. 2). (**E, J, O**) Local dynamic moduli were estimated on the basis of force and deformation magnitude within the ±5-μm area in each profile (grey). Data are mean ± SD. All scale bars are 10 μm. \**p* <0.05, \*\**p*<0.01, two-tailed Student’s *t*-test. N.S., not significant.

Next, we used monastrol, an inhibitor of kinesin-5. Monastrol reduces the affinity of kinesin-5 for the microtubule lattice *in vitro* (Kwok et al., 2006). Although spindles collapsed at a high dose (e.g. 100 μM), bipolarity was maintained when using a relatively low dose (i.e. 10 μM), the efficacy of which was confirmed by the reduced flux velocity (1.0 ± 0.1 μm/min, n = 5; versus 1.7 ± 0.3 μm/min for control, n = 7) (Fig. S4B, C). We found that upon this treatment, the speckle motion profile obtained nearer the spindle pole were similar or slightly suppressed as compared to control (orange in Fig. 5G, H). On the other hand, the profile exhibited a sharper amplitude peak for forces applied near the spindle equator, suggesting enhanced relative filament movement (orange in Fig. 5I). When we increased the monastrol dosage (i.e. 20 μM), spindles shortened to 26.1 ± 4.2 μm (n = 6; versus 37.4 ± 4.8 μm for control) while still maintaining steady length and bipolarity (Fig. S4B, D). Although such small spindles could be analyzed only at two regions, the profiles became much sharper both at the pole and at the equator (red in Fig. 5G, I; Fig. S4E). The estimated local dynamic moduli indicated that the equatorial filament arrays are sensitive to kinesin-5 inhibition, acquiring ~3-fold mechanical compliance upon monastrol treatment (1.4 ± 0.8 kPa at 10 μM; 2.1 ± 0.5 kPa at 20 μM, n = 3 each) (Fig. 5J). The filament array near the spindle pole was less sensitive to this inhibition, but also became compliant upon increasing the dosage (7.5 ± 2.3 kPa at 10 μM; 3.3 ± 1.4 kPa at 20 μM; n = 3 and 5, respectively) (Fig. 5J). Together, these data suggest a broad localization of kinesin-5 across the bipolar structure, and that the activity is required for maintaining spindle length while achieving robust filament couplings near the pole and the equator.

### Dynein contributes to the coupling of microtubule arrays away from the equator

We next focused on dynein, a minus-end directed microtubule motor whose well-characterized function is spindle pole organization (Compton, 1998). An addition of the monoclonal antibody to the dynein light chain (Gaglio et al., 1997) resulted in a barrel-like microtubule array with sprayed poles, the common phenotype of dynein inhibition (Fig. 5K). Because of the unfocused pole, the spindle regions were defined based on the distance solely from the equator such that it nearly matches the region classification of unperturbed spindles (white dotted lines, Fig. 5K). Upon dynein inhibition, we found a noticeable sharpening of the motion amplitude profile at >15 μm away from the equator (Fig. 5L and S4F), consistent with sparse microtubule arrays seen in fluorescence images. Importantly, the difference was also observed at 5-15 μm regions from the equator (Fig. 5M), but was not prominent nearer the equator (<5 μm region) (Fig. 5N). The local stiffness moduli estimated were 1.2 ± 1.5, 0.8 ± 0.3, and 2.6 ± 1.0 kPa (n = 4 each) for regions <5 μm, 5–15 μm, and >15 μm from the equator, among which ≥5 μm regions yielded statistically significant weakening of the structure as compared to control (Fig. 5O). These suggest that dynein contributes to the mechanical coupling of microtubule arrays all along the spindle, except for those around the equator.

## Discussion

Our microneedle-based quantitative micromanipulation with high-resolution fluorescence imaging enabled for directly probing the dynamic changes in position and motility of microtubules in the spindle that respond to applied forces. We found that microtubules at the middle of the spindle half engage in less rigid, more fluid-like mechanical coupling with neighboring filaments and exhibit larger relative movement against perturbing forces, as compared to those around the spindle pole and the equator. Consistent herewith, we discovered that microtubule lattices nearer the spindle pole are extensively slid apart from the equator in response to force altering the pole-to-pole distance, while the arrays of equatorial antiparallel filaments were maintained. From these findings, we propose a model of spindle micromechanics, which is determined by mechanically robust microtubule arrays at the pole and the equator, and more loosely coupled filament arrays at the middle of the spindle half (Fig. 6A).

**Figure 6.**
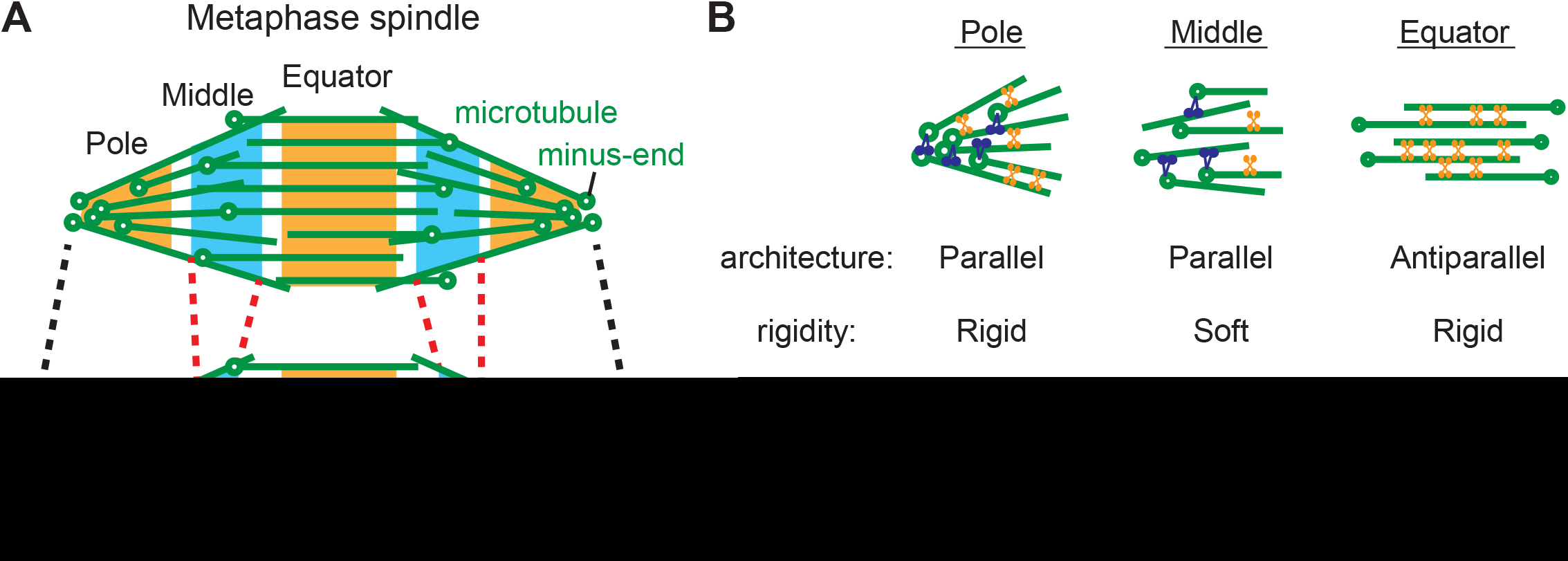
Model for the local mechanical architecture of the spindle. (**A**) Schematic of the metaphase spindle. Open circles, microtubule minus-ends. Short antiparallel microtubules assemble near the equator, while parallel microtubules predominate nearer the pole. The ends of the equatorial and polar microtubule arrays overlap at the middle of the spindle half (highlighted in blue) and form parallel filament arrays, adapting to force associated with spindle-length change (dotted lines). The arrays at the pole and the equator are mechanically more robust and maintain their steady-state architecture against perturbing force (highlighted in orange). (**B**)) Schematic of the molecular interactions involved in spindle micromechanics. Kinesin-5 (orange) localizes across spindle microtubules, pushing and resisting filament sliding to maintain the pole and equatorial dynamics, while its contribution to filament crosslinking at the middle of the spindle half is relatively small. Dynein (blue) localizes at the minus-ends of microtubules and mediates parallel filament interactions at the spindle pole and middle of the spindle half.

Currently, the most advanced models of metazoan spindles describe a steady-state spindle architecture assembled from the collection of short microtubule filaments (Brugues et al., 2012; Burbank et al., 2007; Yang et al., 2008; Yang et al., 2007); however, none explicitly explains the physical nature of the filament interactions and how forces influence their arrangement and motility. . By mechanically perturbing the spindle, we found that there is a predominant fraction of equatorial microtubules whose mechanical coupling to the spindle pole is considerably weak, and thus, their dynamics is insensitive to a force that pulls the two poles apart. On the other hand, microtubule arrays nearer the spindle pole engage in more rigid mechanical connection and their movement is tightly coupled to spindle-length change. Given the shortness of the filaments, the microtubule array at the middle of each spindle half is likely formed by overlapping ends of equatorial antiparallel filaments and polar parallel filaments (blue highlighted area in Fig. 6A). Our data show that this predominantly parallel filaments array has less mechanical resistance than other spindle location and can adapt to force that alters the pole-to-pole distance. Further, microtubules growing from the opposite spindle pole engage in nearly equivalent mechanical coupling with neighboring filaments, masking their structural polarity. These filament-coupling features, which are relatively weak, spatially heterogeneous, and polarity-independent, enable the spindle to transmit and respond to force in a manner that is distinct from the long-standing model of spindle assembly, in which the arrays of equatorial antiparallel microtubules connect two spindle poles and balance the pole-to-pole distance (Inoue and Salmon, 1995; Mitchison and Salmon, 2001; Scholey et al., 2003).

Our results also suggest how the robust and adaptable nature of the microtubule arrays emerges from motor protein mechanics. At the spindle equator, the major protein that maintains the filament dynamics is likely kinesin-5, as our data showed the sensitivity of the equatorial mechanics to chemical inhibitor. When kinesin-5 was inhibited, filament sliding slowed down, most likely because of a lower number of force generators acting against a constant load, and further, the filaments more easily slid apart against the force perturbing their motility. Unlike other kinesins, such as vesicle transporting kinesin-1, *Xenopus* kinesin-5 can maintain a stable association with the lattices of microtubules and generate substantial braking force against fast filament sliding (Shimamoto et al., 2015; Valentine et al., 2006). This resistive motor force is additive (Shimamoto et al., 2015) and can thus accumulate across the filament overlap of several microns at which many kinesin-5 molecules localize (Kashina et al., 1996). Because of these motor properties, each kinesin-5 molecule should experience only a subtle force fluctuation that is insufficient to perturb the intrinsic enzymatic cycle, thus maintaining the speed of antiparallel filament sliding and preventing excess filament translocation against perturbing forces (“Equator” in Fig. 6B). Our inhibition assays suggested that, around the spindle pole, kinesin-5 also crosslinks parallel microtubules and enhances filament coupling, as predicted from its substantial pole localization (Sawin et al., 1992) and reconstitution assays (Kapitein et al., 2005; Shimamoto et al., 2015). Filament coupling at the pole also depended on dynein, in agreement with previous studies (Gaglio et al., 1997; Tan et al., 2018). Thus, kinesin-5 and dynein may act together to make the rigid filament coupling when microtubules reach the pole (“Pole” in Fig. 6B). In contrast to these robust structures, however, those at the middle of the spindle half appeared much more compliant. Our findings suggest that within this filament array where predominant microtubules run in parallel, kinesin-5 crosslinking activity is largely suppressed (“Middle” in Fig. 6B). On the other hand, dynein plays a prominent role, likely via its minus-end accumulation and lateral interaction with adjacent filaments (Hueschen et al., 2017; Tan et al., 2018) (“Middle” in Fig. 6B). Based on our measurement of local spindle stiffness (~1,000 pN/μm_2_) and the density of microtubules (~100/μm_2_), the linkage could generate ~10 pN force per filament, an amount 2-to 5-fold larger than the stall force of single dynein molecules walking along a single microtubule (Gennerich et al., 2007; McKenney et al., 2010; Torisawa et al., 2014). The parallel filament interaction could also be mediated by non-motor microtubule-associated proteins such as the augmin complex, which caps the filament minus-ends and promotes microtubule branching (Goshima et al., 2008). This might be the basis of the mechanical resistance that was observed upon dynein inhibition. The viscous, fluid-like property identified nearer the equator reflects the dynamicity of crosslinkers that allows for filament re-arrangements, whereas the elastic property nearer the pole suggests static, persistent cross-bridges. Our work thus predicts a rich micromechanics underlying parallel microtubules, which is less appreciated but as important as that in antiparallel filaments.

In the spindle, chromosomes are captured by another subset of parallel microtubules, called kinetochore microtubules or k-fibers, which run across the spindle half and pull the chromatids toward the poles. Although this filament fraction was minor in our present study (<10% of total filament number), their length must also be coupled to spindle length in order to prevent chromosomes from detaching. A recent electron tomography study revealed that kinetochore microtubules of *C. elegans* spindles are not continuous, but rather, their minus-ends are terminated midway and embedded within a network of short microtubules assembled around the pole (Redemann et al., 2017). In addition to the previously identified mechanism regulating filament depolymerization at the fiber ends (Dumont and Mitchison, 2009a; Skibbens and Salmon, 1997), we predict that the mechanical linkage between the filaments is compliant and can adapt to force. Capturing the motility of individual microtubules within this thick filament bundle would be an important next challenge to build a complete model of spindle micromechanics.

Our findings on spindle’s mechanical heterogeneity suggest functional advantages as it allows for maintaining the equatorial spindle dynamics while controlling the pole-to-pole distance. The equatorial microtubule arrays serve as structural scaffolds for physical stretching of kinetochores (Elting et al., 2017) and for biochemical signaling of cytokinetic furrow positioning (Oegema and Mitchison, 1997). We anticipate that the mechanically distinct local microtubule arrays maintain these spindle functions, ensuring the robustness of chromosome segregation while adapting to perturbation for error-free cell division.

## Materials and Methods

### Spindle assembly

Metaphase spindles were assembled in extracts prepared from unfertilized *X. laevis* eggs according to a well-established method (Desai et al., 1999). Freshly prepared, cytostatic factor-arrested extracts (30 μl per reaction) were first supplemented with demembranated *X. laevis* sperm nuclei (400 nuclei/μl) and released into interphase by addition of Ca_2+_ at a final concentration of 0.4 mM. Following 90-min incubation at 18°C, reactions were diluted with the equal volume of fresh extracts and cycled back into metaphase. Following 50-min incubation, X-rhodamine-labeled tubulin and Alexa 488-labelled tubulin, prepared according to a previously described method (Hyman et al., 1991), were added to extracts at a final concentration of 20 nM and 800 nM, respectively. SYTOX Green (S7020, Invitrogen) was also added to extracts at a final concentration of 250 nM for chromosome imaging. Experiments were performed 60–150 min from the start of spindle assembly, during which no noticeable changes in spindle mechanics and overall morphology were observed.

### Microneedles

Microneedles were prepared by pulling glass rods (G1000, Narishige) using a capillary puller (PC-10, Narishige) followed by processing of their tips using a microforge (MF-100, World Precision Instruments) (Shimamoto and Kapoor, 2012). For precise control over its position and movement in viscous egg extracts while probing spindle force with sufficient sensitivity, the tip of each force-calibrated microneedle was made ~1–2 μm in diameter and ~100–300 μm in length, which yielded a stiffness of 0.3–0.5 nN/μm as determined by the cross-calibration method (Shimamoto and Kapoor, 2012). The tips of microneedles used in spindle-stretching experiments were made >100-times stiffer, with ~2-μm diameter and ~50-μm length.

### Microscopy

Spindle micromanipulation was carried out in an inverted light microscope (Ti-E, Nikon) equipped with a pair of three-axis hydraulic micromanipulators (MHW-3, Narishige), a 100× objective (Apo TIRF, 1.49NA, Nikon), an objective scanner (PIFOC, Physik Instrumente), a motorized sample stage (MS-2000, Applied Scientific Instruments), a spinning-disk confocal unit (CSU-X1, Yokogawa), and an sCMOS camera (Neo 4.0, Andor). Two excitation lasers (488 nm and 561 nm, 40 mW, OBSI, Coherent) were merged using an in house-built laser combiner and were introduced into the confocal unit via an optical fiber (Yokogawa). The microscope and imaging instruments were wired to a computer and controlled using image acquisition software (NIS-Elements, ver. 4.50, Nikon).

For oscillatory perturbation experiments, a single-microneedle setup was used. First, 4 μL of a cycling extract containing pre-assembled spindles was placed in an open experimental chamber, which was assembled from a coverslip (Matsunami) and a rubber plate with 10-mm central aperture, and covered with mineral oil (M-5310, Sigma). Under a confocal microscope, a bipolar spindle of typical size and shape was selected under low illumination conditions, and the tip of a force-calibrated microneedle was inserted into the region of interest within the spindle. The microneedle tip was brought down to 1–2 μm above the coverslip while maintaining nearly vertical approaching angle (>80° with respect to the horizontal plane). The microneedle base was held by a translational piezo actuator (P-841.20, Physik Instrumente) whose displacement was controlled by a voltage signal generated in an in house-written LabView program (National Instruments) and sent via a closed-loop piezo driver (E-665, Physik Instrumente). Experiments were performed using a sinusoidal force (frequency: 0.1 Hz) applied by moving the microneedle tip parallel to the spindle pole-to-pole axis. The amount of force applied was estimated based on the displacement of the microneedle tip from its equilibrium position, which was determined using time-lapse imaging of the tip and piezo sensor reading. Time-lapse images were acquired at a single confocal plane (~5 μm from the coverslip surface) with pre-optimized image acquisition settings (interval: 1 s; exposure time: 200 ms for 488 nm and 500 ms for 561 nm) that fulfilled the following requirements: 1) photo-damage and photobleaching were minimal, and 2) individual tubulin speckles could be tracked across the time-lapse sequence.

For microrheology analysis, a piezo-based nano-positioning stage (Nano-LP200, Mad City Lab) was mounted onto the motorized sample stage and moved in a sinusoidal manner at a fixed frequency (0.02–4 Hz) and amplitude (0.7–1.0 μm) along the pole-to-pole axis of the spindle. The base of the force-calibrated microneedle was held at a fixed position throughout the measurement while its tip was inserted into the spindle. Bright-field images were acquired at the sampling rate 50-times the frequency of the oscillatory force input for measurements at ≤0.2 Hz (e.g. 1,000 ms interval for 0.02 Hz input), and at a fixed 40-ms interval for measurements at >0.2 Hz. The magnitude of applied force was monitored based on the microneedle tip’s displacement. The amount of spindle deformation was estimated by the relative displacement of the microneedle tip and the spindle, whose position change was monitored by tracking the center of a tracer microbead (LB30, Sigma) immobilized onto the coverslip surface.

For stretching perturbation experiments, the microscope setup as described above was used, but with a dual-microneedle setup (Takagi et al., 2014). In each experiment, a single bipolar spindle of typical size and shape was captured by inserting the tips of the microneedles near the opposite spindle poles (<5 μm from the structure’s edge). One microneedle was used to pin down the spindle while the other microneedle was used to stretch the bipolar structure at a constant velocity (100 nm/s). The stretching motion of the microneedle was controlled by either manual steering of the micromanipulator or using the piezo actuator attached to the microneedle base. Time-lapse images were acquired in a single confocal plane at 3-s intervals while switching two excitation lasers with <200 ms time delay (exposure time: 400 ms for 488 nm and 300 ms for 561 nm). Spindles displaying no visible elongation in response to the micromanipulation (<10% of the original steady-state length) usually associated with pole disorganization and were excluded from subsequent analyses.

### Molecular perturbation

AMPPNP (Sigma) was used at a final concentration of 1.5 mM after adjustment of the pH to ~7.7 with potassium hydroxide. Monastrol (M8515, Sigma) was used at a final concentration of 10 or 20 μM in the presence of 0.5% DMSO. Monoclonal antibody to the dynein light chain (D5167, Sigma) was first dialyzed against a buffer comprised of 50 mM potassium glutamate and 0.5 mM MgCl2 using a centrifugal membrane filter (Amicon Ultra, Millipore) and then added to extracts at a final concentration of 1 mg/ml (Heald et al., 1997). Reagents were prepared as 50–100× working stocks in CSF-XB (10 mM K-Hepes, pH 7.7, 1 mM Mg_2+_, 1 mM EGTA, 150 mM KCl, 50 mM sucrose) and were added to extracts containing pre-assembled spindles.

### Data analysis

Spindle length was measured on the basis of fluorescence images of Alexa 488-tubulin. In each image of a time-lapse sequence, a line-scan was performed along the pole-to-pole axis of the spindle. The edges of the line-scan profile were detected on the basis of an intensity threshold that was set at 25% of the maximal spindle signal intensity. The distance between the two edge positions was defined as the spindle length.

Speckle motion was analyzed on the basis of fluorescence images of X-rhodamine-tubulin. The entire time-lapse image stack from each experiment was first low-pass filtered (pixel width: 2 × 2) in the NIS-Elements software and then loaded in the Particle Track and Analysis plugin (https://github.com/arayoshipta/PTA2) in ImageJ. Speckles were detected on the basis of the total intensity and size of fluorescence spots that exceeded fixed threshold values, and then, their intensity profiles were each fit to a two-dimensional Gaussian function for calculating the centroid position. The speckles that were detected in each image were then linked across the time-lapse sequence, with fixed linking parameters. After visual inspection of representative speckle trajectories, the motion of each speckle was analyzed as follows:

1. *Oscillatory perturbation experiments*. The time recording of each speckle displacement along the long spindle axis (*x_L_*) was fit to a sinusoidal function, which was given by *x_L_* (*t*) = *A* sin(*ωt* + *θ*) + *Bt* + *C*. Here, *A* is the amplitude of speckle motion, *ω* is the angular frequency corresponding to the input force sinusoid (0.1 Hz, *ω* ~ 0.63 rad/s), *θ* is the oscillatory phase, and *t* is the time elapsed from the onset of force application. The variable *B* is to compensate translational movements of speckles associated with the poleward flux; a plus or minus sign was used to assign the polarity of each microtubule filament. The variable *C* is to correct initial position offset. Fitting was conducted in Origin 2016 (Origin Lab) and data that yielded an *R^2^* value above 0.25 were used for subsequent analyses. The profile of speckle motion amplitude along the long and short spindle axes was generated on the basis of data of speckles whose initial tracking point was included within a ROI. The ROI was drawn along each spindle axis (width: ± 2.5 μm). After the removal of outliers (defined as speckles exceeding the peak amplitude of the speckle closest to the force application point) followed by smoothing of the data plots (using 5-μm moving average filter) and offset correction (subtracting the translational drift of the entire structure, ~0.1 μm typical), each profile was aligned at the local maximum within a ±5-μm region from the initial microneedle position, and profiles were averaged over multiple spindle samples.
2. *Spindle stretching experiments*. For individual speckle trajectories acquired during the steady lengthening phase (see the Results section), the instantaneous velocity of the speckle was calculated by dividing the contour length of each trajectory by the period over which the speckle was tracked. Speckles that could be tracked over ≥10 s (≥3 successive time-lapse frames) were used for subsequent analyses. The velocity data were plotted in a single imaging plane based on the position relative to the spindle equator, which was calculated using the initial tracking point of each speckle and the length and width of the spindle at the corresponding time point.

The magnitude of the force (*F*) applied was estimated on the basis of the pre-calibrated microneedle tip stiffness (*k_M_*) and its displacement from the equilibrium position (*Δx_M_*) according to Hooke’s law, which is given by *F = k_M_ Δx_M_*.

For microrheology analysis, time-dependent changes of the force and spindle deformation were each fit to a sinusoidal function, which was given by *A(t) = A_0_* sin (*ωt + θ*), and used to determine the amplitude and phase. The effective stiffness was determined by the ratio of the force to the deformation amplitude. The phase shift was determined by the difference between the two oscillatory phases.

Local spindle region (i.e. pole, equator, and middle) was classified on the basis of the distance from either the structure’s center or its outer edge, measured along the pole-to-pole axis of the spindle. The pole region was defined as the area within 5 μm from the structure’s outer edge. The equator region was defined as the area within 5 μm from the structure’s center. The middle region was defined as the area between the two above regions. Based on these definitions, reduced-sized spindles at 20 μM monastrol allowed for data acquisition at only the equatorial and pole regions. For spindles with dynein inhibition, the regions were defined solely on the basis of the distance from the structure’s center (<5 μm, 5–15 μm, and >15 μm) because of defocused spindle poles.

The dynamic modulus of the spindle was estimated as follows. First, we measured local deformation of microtubule arrays that developed within a 5 × 5-μm area around the point of force application, analyzed on the basis of relative speckle movement in the *x-y* plane (i.e. the imaging plane; *x*, long spindle axis; *y*, short spindle axis) and assuming that the deformation was even across the *z*-axis (i.e. over the entire spindle width). Below, we use the following indices for each coordinate: *x* = 3, *y* = 1, and *z* = 2. The local deformation was described by strain tensor *E_ij_*, where *i* and *j* are the directions of strain and normal plane, respectively. On the other hand, the stress developed within the area was described by stress tensor *p_ij_*, where indices are identical to those of the strain tensor. These two tensors can be related by the dynamic modulus tensor *C_ijkl_*, as *P_ij_* = *C_ijkl_ E_kl_*. Here, *P_ij_* is the symmetric tensor and thus, *P_12_ = P_21_, P_13_ = P_31_*, and *P_32_ = P_23_*, and similarly for *E_kl_*. Also, *C_ijkl_ = C_jikl_ =C_ijkl_ = C_klij_* We also considered transverse isotropy of the bipolar spindle, where the y-z plane is the isotropic plane, and thus, *C_55_ = C_44_* and *C_22_ = C_11_* as Voigt index. Under these assumptions, the longitudinal stress *P_33_* can be described as: *P_33_ = C_13_E_11_ + C_13_ E_22_ + C_33_ E_33_*. The transverse normal strain we measured experimentally was negligibly small and therefore, *E_11_* and *E_22_* were omitted. The longitudinal strain *E_33_* was estimated by the difference in displacements at the peak and edge positions (averaged over values at the two edges) within the defined 5 × 5-μm area. On the other hand, the longitudinal stress *P_33_* was defined as the magnitude of force measured by the microneedle probe and the area normal to the force application vector. The lateral dynamic modulus was estimated under the same assumptions and was given by *P_31_ = P_32_ = C_44_ E_31_*.

The statistical tests were performed in Origin Pro 9 (Origin Lab Corp) on the basis of two-tailed Student’s *t*-test.

## Supplementary information

Supporting data of this manuscript are provided as a separate PDF file. The PDF file includes Supplementary Figures 1–4, and captions of the figures.

## Acknowledgements

We thank Dr. Akatsuki Kimura, Dr. Tarun Kapoor, and Dr. Shin’ichi Ishiwata for critical reading of the manuscript, and members of Shimamoto Lab for valuable comments and suggestions. This work was supported by JSPS KAKENHI 16H06166, 17K19362 (to Y.S) and JSPS Postdoctoral Fellowship (to J.T.).

## Author contributions

J.T. and Y.S. designed assays, performed experiments, analyzed data, discussed results, and wrote the manuscript.

